# Immunosuppression Drugs Exhibit Differential Effects on Endothelial Cell Function

**DOI:** 10.1101/2024.10.31.620858

**Authors:** Aly Elezaby, Ryan Dexheimer, David Wu, Sze Yu Chan, Ian Y. Chen, Nazish Sayed, Karim Sallam

**Author notes:** Corresponding Author: Karim Sallam, MD.

## Abstract

Immunosuppressive medications are widely used to treat patients with neoplasms, autoimmune conditions, and solid organ transplants. Prior studies indicate that immunosuppression drugs can cause adverse vascular remodeling. Given the systemic effects of the drugs, elucidating cell-type specific drug-effects has been challenging. We utilized induced pluripotent stem-cell derived endothelial cells to investigate the role of widely used immunosuppression drugs on endothelial function. We found that among immunosuppression agents, sirolimus reduced basic endothelial cell functions including cell migration, proliferation, acetylated LDL uptake, and angiogenesis properties; while tacrolimus only reduced nitric oxide release. This model allows for investigation of differential effect of immunosuppression drugs on endothelial function that can elucidate mechanisms contributing to adverse vascular profiles observed clinically.

## Introduction

Immunosuppressant agents, including mTOR (mammalian target of rapamycin) inhibitors and calcineurin inhibitors, are essential in preventing organ rejection following transplantation^1^. However, their use is associated with potential side effects, specifically impacting the vascular system with long-term complications^2^. While calcineurin inhibitors are associated with an increased risk of hypertension, mTOR inhibitors, such as sirolimus, are associated with a lower incidence of hypertension^3^. However, mTOR inhibitors have been linked to the development of peripheral edema and lymphedema^4^. Furthermore, while mTOR inhibitor use increases serum lipids^5^, they paradoxically are associated with a lower risk of atherosclerosis^6^.

In heart transplantation, immunosuppressants have variable effects on the attenuation of cardiac allograft vasculopathy (CAV), a condition marked by pathologic vascular remodeling and fibrosis and a major driver of graft dysfunction and mortality^7^. Calcineurin inhibitors are largely associated with an increased risk of CAV, while mTOR inhibitors attenuate this risk^8^. Even though immunosuppressants are known to cause endothelial injury, inflammation, and dysregulation of vascular smooth muscle cell proliferation, the precise mechanisms by which immunosuppressants contribute to transplant vasculopathy are not yet fully understood^9^. Thus, elucidating the cellular subtypes and the cell-specific changes that drives abnormal vascular phenotype can help highlight therapeutic targets to attenuate pathologic vascular remodeling and minimizing cardiovascular complications in transplant recipients.

Our aim here is to investigate the direct role of immunosuppressive agents on the endothelium. Human induced-pluripotent stem cell-derived endothelial cells (iPSC-EC) were used to investigate the effects of calcineurin inhibitors (tacrolimus) and mTOR inhibitors (sirolimus) on critical components of vascular remodeling.

## Methods

### Endothelial Cell Differentiation

iPSCs used in this study were provided by the Stanford Cardiovascular Institute Biobank. We cultured these iPSCs in StemMACS™ iPS-Brew media until cells have reached 85% confluence. Differentiation was started with RPMI-B27 minus insulin media (Life Technologies) with 6uM CHIR99021 (GSK3 inhibitor) for 2 days to drive cells down the mesodermal lineage. From days 2 to 4, cells are cultured in RPMI-B27 minus insulin media with 2uM CHIR99021 media. From days 4 to 12, cells are cultured in EGM™-2 Endothelial Cell Growth Medium-2 (Lonza CC-3162) supplemented with 50 ng/mL vascular endothelial growth factor (PeproTech), 10 uM SB 431542 (Tocris), and 20 ng/mL fibroblast growth factor 2 (PeproTech) where media is changed on days 4, 6, 8, and 10. On day 12, cells are dissociated with TrypLE for 6-8 minutes and non-endothelial cells were separated through MAC sorting with CD144 MicroBeads (Miltenyi Biotec). Purified endothelial cells (passage 0) were seeded onto 10cm dishes coated with .2% gelatin and cultured in EGM2 medium with 10uM SB 431542. To passage iPSC-ECs, cells were dissociated with TrypLE for 5 minutes and transferred to plates coated with .2% gelatin and cultured in EGM2 media without supplements for downstream experiments (passage 1). The iPSC-ECs used in this study were cultured until passage 2.

### Angiogenesis (Tube Formation Assay)

The angiogenesis assay was adapted from a previously established protocol^10^. Endothelial cells (ECs) derived from iPSCs, or iPSC-ECs, were treated with vehicle (control), tacrolimus (1uM), and sirolimus (50nM) on a 6-well plate for 5 days and seeded at 5 x 10^4^ cells in complete EGM2™ media (Lonza Bioscience, CC-3156 and CC-4176) on Basement Membrane Matrix (Fisher Scientific, CB-40234A)-coated 24-well plates. Cells were incubated at 37°C, 5% CO_2_, and 20% O_2_ for 8 hours and underwent brightfield imaging at 10x magnification on a Keyence BZ-X800. Images were analyzed using a well-established angiogenesis analyzer plug-in on ImageJ to quantify total number of branches, branch length, etc^11^. On ImageJ, the raw image files were converted to RGB color and then analyzed using the “HUVEC Phase Contrast” option under the “Network Analysis Menu.” The results were exported from ImageJ and analyzed using One-Way ANOVA on GraphPad Prism 10.

### Scratch-Wound Assay

The scratch-wound assay was adapted from a previously established protocol^12^. ECs were cultured on a 12-well plate until 95-100% confluent and treated for 5 days with vehicle (control), tacrolimus (1uM), or sirolimus (50nM). A single line, approximately 500 um wide, was scratched in each well. After washing, fresh complete EGM2™ was added to the cells and wells underwent serial brightfield imaging of the scratch at 10x magnification on the Keyence BZ-X800 at 0 hours and at 24 hours (after incubation at 37°C, 5% CO_2_, and 20% O_2_). Images were analyzed manually using ImageJ to measure cell-free scratch area. The speed of cell migration was determined from the change in the width of the empty space from 0 hours to 24 hours divided by 24 hours. For each well, the scratch closure percentage was calculated by dividing the change in empty space over the 24 hours by the initial scratched area. These values were exported to GraphPad Prism 10 and a One-Way ANOVA was performed to determine statistical significance.

### Nitric Oxide Assay

ECs were cultured at 95-100% confluency on a 6-well plate and treated for 4 hours with vehicle (control), tacrolimus (1uM), and sirolimus (50nM). After washing, cells were incubated in Opti-MEM reduced serum medium (ThermoFisher Scientific, 31985062) at 37°C, 5% CO_2_, and 20% O_2_ for 1 hr. The nitric oxide assay was performed per Greiss Reagent Kit protocol for nitrite determination (ThermoFisher Scientific, G7921). Experimental samples were measured in triplicates and standard curve was measured in duplicates. In a clear 96-well plate, 150 uL of the experimental samples (in triplicates) and standard solutions (in duplicates) were added along with 20 uL of prepared Greiss Reagent, incubated at room temperature for 30 minutes and absorbance measured using a Cytation5 (BioTek) to determine nitric oxide concentrations based on standard curve. These values were exported to GraphPad Prism 10 and analyzed by one-way ANOVA to determine statistical significance between treatment groups.

### Low-Density Lipoprotein (LDL) Fluorescence Assay

ECs were cultured in a 6-well plate and treated with vehicle (control), tacrolimus (1uM), and sirolimus (50nM) for 5 days. The cells were incubated at 37°C, 5% CO_2_, and 20% O_2_ for 5 minutes and then the cells were gently pipetted off the wells. The TrypLE-cell suspension mix was added at a 1:1 ratio to complete EGM2™ media in a 15 mL confocal tube. The cells were centrifuged for 5 minutes at 300 rpm. EGM2™ was used to resuspend the cells and the number of ECs was calculated using a Countess 3 automatic cell counter (Invitrogen). In 8-well chamber slides, coated with Matrigel™ (Fisher Scientific, CB-40230C), ECs were seeded at 6 x 10^3^ cells in complete EGM2 media and incubated at 37°C, 5% CO_2_, and 20% O_2_ for 3 hours, and subsequently serum-starved overnight. Acetylated Low-Density Lipoprotein Alexa Fluor™ 594 conjugate (ThermoFisher Scientific, L35353) was added directly to each well and incubated at 37°C for 4 hours. After incubation, the cells were washed with PBS containing calcium and magnesium (ThermoFisher Scientific, 14040133). A cover slip was placed on the slide and immediately imaged. Fluorescent imaging was performed at 20x using a Texas Red filter on the Keyence BZ-X800. Corrected total cell fluorescence was performed via ImageJ. For each image, intracellular fluorescence was measured, corrected for cell area and background fluorescence. These values were analyzed via one-way ANOVA on GraphPad Prism 10.

### Seahorse Assay (XF Real-Time ATP Rate Assay)

The Seahorse ATP Rate Assay was conducted per manufacturer’s protocol (Agilent Technologies). ECs were cultured on a XFe96 Seahorse microplate (Agilent Technologies, 103794-100) or XFe24 Seahorse microplate (Agilent Technologies, 100882-004) until 95-100% confluent and treated for 5 days prior to performing the Seahorse assay. On the day of the experiment, cells were incubated in Seahorse XF DMEM medium (Agilent Technologies, 103575-100) supplemented with 1mM sodium pyruvate (Agilent Technologies, 103578-100), 10mM glucose (Agilent Technologies, 103577-100), and 2mM L-glutamine (Agilent Technologies, 103579-100). Per ATP Rate Assay protocol, oxygen consumption rate (OCR) and extracellular acidification rate (ECAR) were measured in cells treated with 1.5uM oligomycin (via Port A) followed by 0.5uM antimycin A and 0.5uM rotenone (via Port B) (Agilent Technologies, 103592-100). At the end of the Seahorse experiment, cell count was completed using NucBlue™ Live ReadyProbes™ (ThermoFisher Scientific, R37605) using a BioTek Cytation5 to normalize the ATP rate values. Statistical analysis was performed with one-way ANOVA in GraphPad Prism 10.

### Statistical Analysis

Values are presented as mean ± S.D. relative to the average value for the control group unless otherwise stated. In experiments with two groups, group differences were assessed by Student’s unpaired two-tailed t-test. In experiments with more than two groups, one-way ANOVA with Tukey’s multiple comparison test was used. Statistical significance was established at *p* < 0.05 (Graph Pad Prism).

## Results

We examined the effects of immunosuppressant agents on endothelial cells derived from iPSCs from three patient control lines using established monolayer protocols. iPSC-ECs treated with tacrolimus and sirolimus revealed alterations in endothelial cell function with significant differences between the drug classes.

We conducted a scratch assay to examine the differential effects of tacrolimus and sirolimus on endothelial cell migration and wound healing **(Figure 1)**. The assay implicates a manual disruption of endothelial monolayer “scratch” and then assess the size of the cell-free zone in the monolayer 24 hours later. iPSC-ECs treated with sirolimus showed 68% decrease in cell migration compared to controls, whereas tacrolimus treatment did not differ from control with respect to cell migration. In addition, sirolimus resulted in a 72% decrease in scratch closure compared to the controls. Tacrolimus treatment, on the other hand, did not result in any significant change in scratch closure compared to the control group.

**Fig 1.**
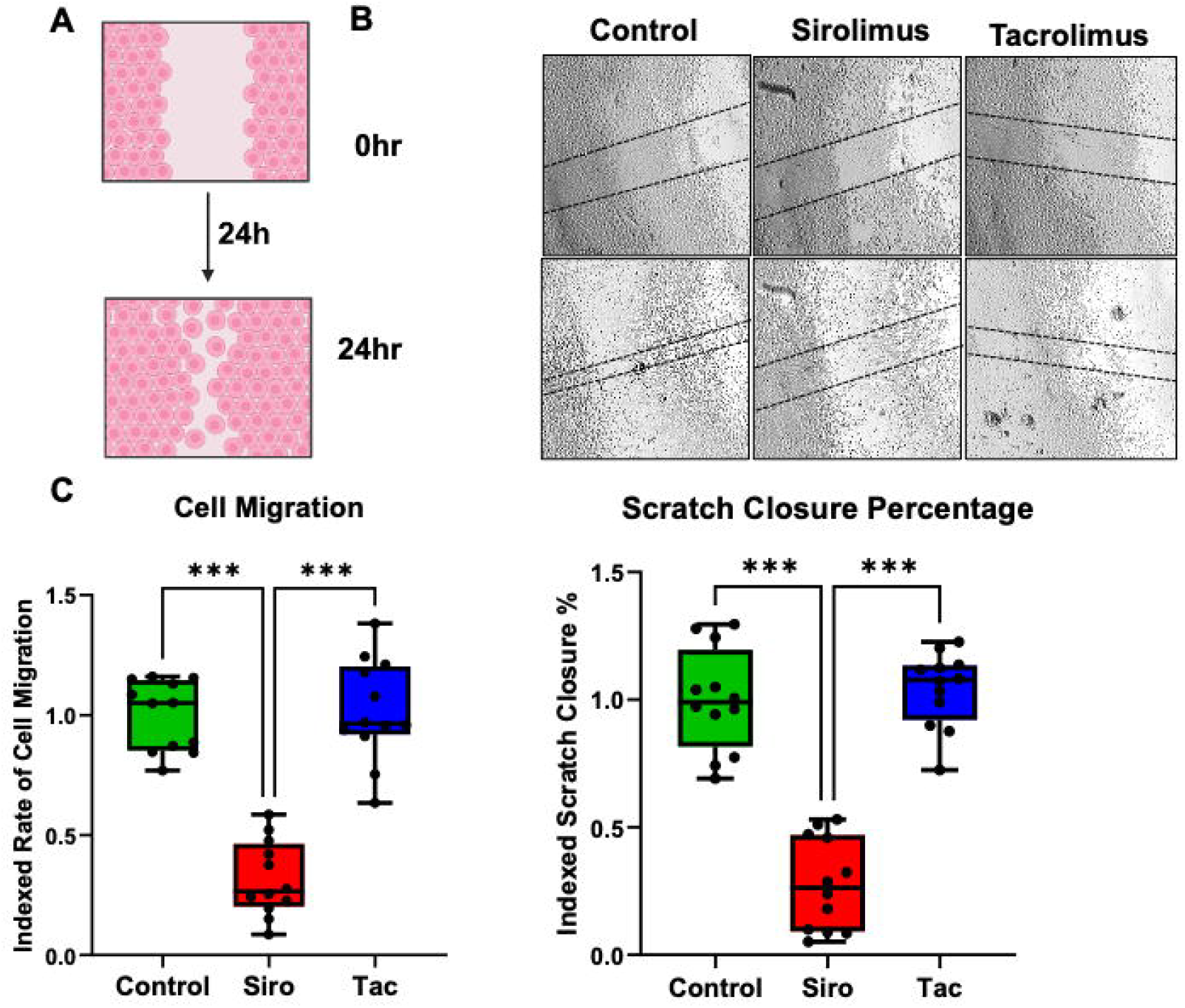
Sirolimus-treated endothelial cells exhibit decreased cell migration and scratch closure. (A) Schematic of scratch assay. (B) Representative images of scratch closure at times 0 and 24 hours in vehicle-treated control cells, sirolimus-treated cells, and tacrolimus-treated cells. (C) Indexed rate of cell migration and scratch closure percentage in cells treated with sirolimus (Siro), tacrolimus (Tac), or vehicle (Control).

Prior studies have indicated that both calcineurin and mTOR inhibitors are associated with dyslipidemia^3,13,14^. To determine the effect of immunosuppression agents on LDL uptake by iPSC-ECs, we examined LDL levels in cells after treatment with sirolimus or tacrolimus **(Figure 2)**. Sirolimus resulted in a significant reduction in acetylated LDL uptake compared to control or tacrolimus treatment. Tacrolimus and control groups LDL uptake did not differ significantly. The reduction in LDL uptake may explain the lower risk of atherosclerosis associated with sirolimus despite the drug being associated with dyslipidemia^15^.

**Fig 2.**
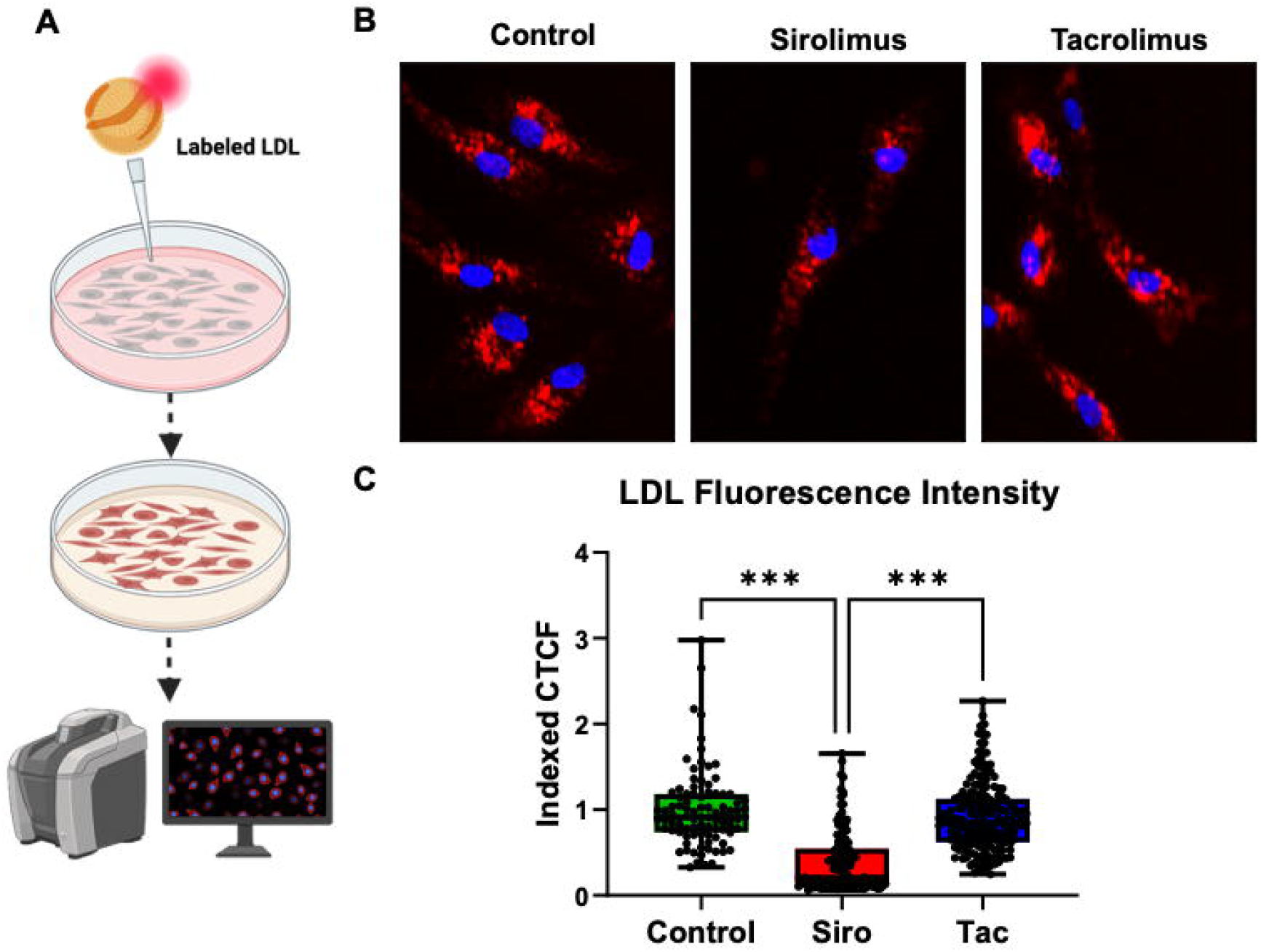
Sirolimus-treated endothelial cells have decreased LDL uptake. (A) Schematic of acetylated LDL uptake assay in cells treated with sirolimus, tacrolimus or vehicle control. (B) Representative image of fluorescent acetylated LDL uptake in iPSC-ECs treated with sirolimus, tacrolimus or vehicle control. (C) Quantification of LDL fluorescence intensity in iPSC-ECs treated with sirolimus (Siro), tacrolimus (Tac), or vehicle (Control).

mTOR inhibitors have been shown to inhibit pathological angiogenesis (in the context of malignancy) as well as physiological angiogenesis (occurring with wound healing and development)^16^. To examine the relative effects of sirolimus and tacrolimus on angiogenesis, we performed a tube formation assay. This in vitro assay surveys the natural tendency of endothelial cells in culture to form tube networks which is disrupted by endothelial dysfunction. We found that sirolimus-treated iPSC-ECs had limited branching capability, with a decrease in branch numbers, network length and branch length relative to controls. Sirolimus also exhibited a higher number of isolated segments and isolated total segment length compared to control, consistent with poor tube formation. Conversely, cells treated with tacrolimus showed no difference in branch number, branch length, number of isolated segments, or isolated segment length branch length compared to controls **(Figure 3)**. These findings are indicative of reduced angiogenesis with sirolimus but not tacrolimus treatment.

**Fig 3.**
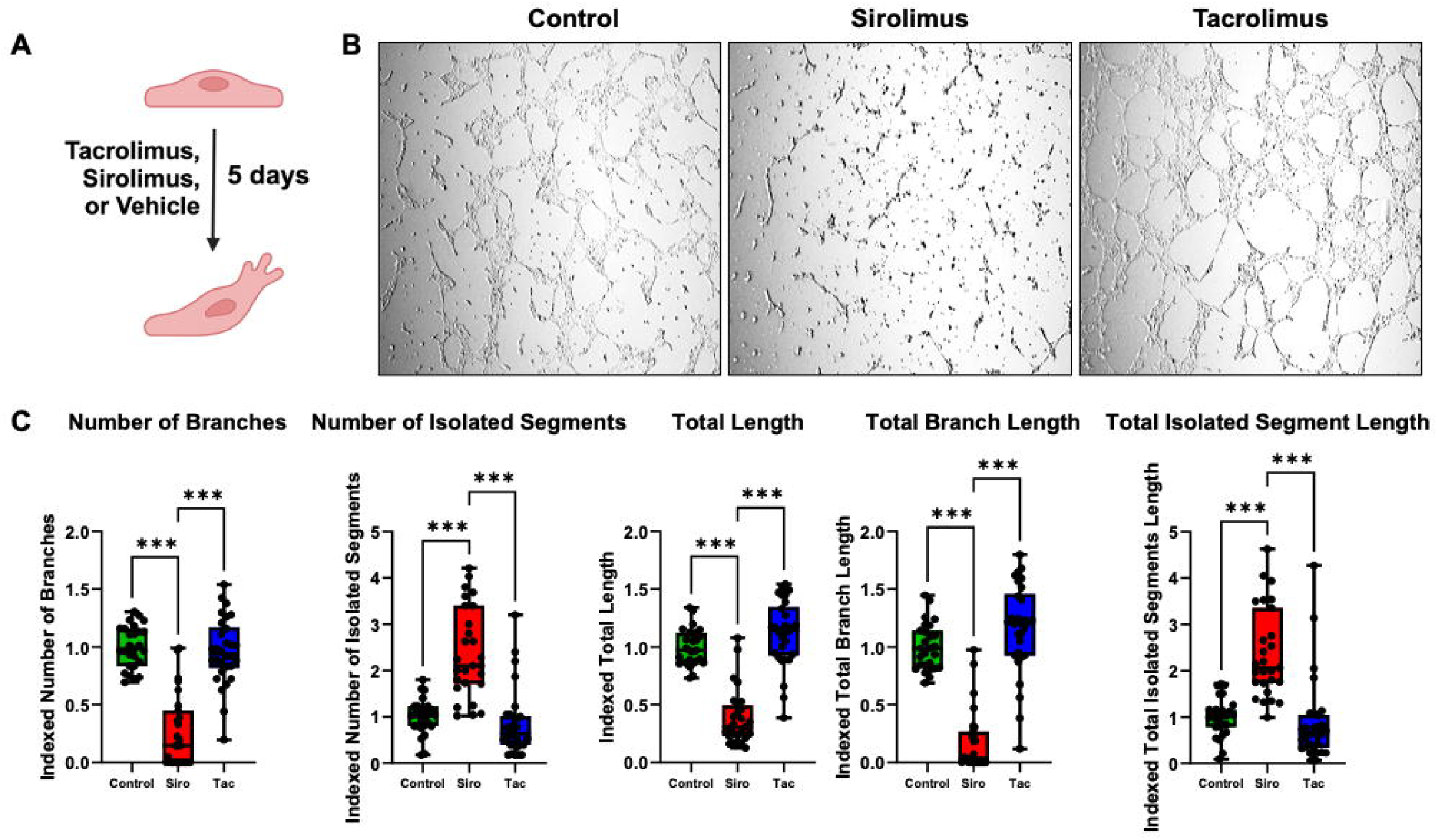
Sirolimus-treated iPSC-ECs have diminished angiogenic potential. (A) Schematic of tube formation assay in iPSC-ECs treated with sirolimus, tacrolimus or vehicle control for 5 days. (B) Representative brightfield microscopy images showing differences in branch formation in iPSC-ECs treated with sirolimus, tacrolimus or vehicle control. (C) iPSC-ECs treated with sirolimus have a decrease in number of branches, number of segments, total length, branch length and isolated segment length.

Endothelial cells produce and release nitric oxide to induce vasodilation. Additionally, nitric oxide produced in the vascular lumen prevents platelet aggregation and leukocyte adhesion, thereby decreasing atherogenesis^17^. The use of calcineurin inhibitors and mTOR inhibitors is associated with a differential effect on atherosclerosis^6,18^. We sought to determine the relative effects of sirolimus and tacrolimus on nitric oxide release from endothelial cells. Both sirolimus and tacrolimus treatment reduced nitric oxide production. Sirolimus decreased nitric oxide release from iPSC-ECs by 47% and tacrolimus by 28% compared to controls **(Figure 4)**. This is consistent with impaired nitric oxide release in response to both immunosuppression agents.

**Fig 4.**
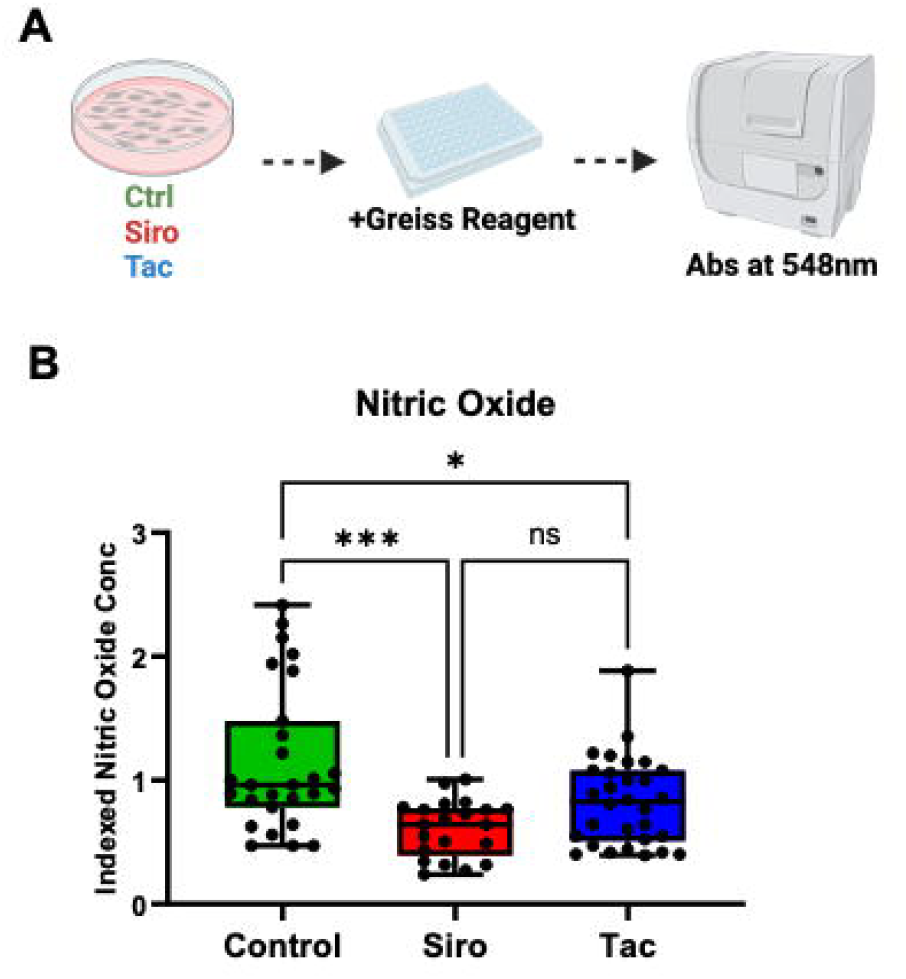
Sirolimus and tacrolimus decrease nitric oxide production. (A) schematic of nitric oxide production assay from iPSC-ECs treated with sirolimus, tacrolimus or vehicle control. (B) iPSC-ECs treated with sirolimus and tacrolimus have a decrease in nitric oxide production relative to control.

Endothelial cell function has been linked to mitochondrial energetics^19^, and immunosuppressive agents can exert direct effects on mitochondria to impact metabolism and energetics^20,21^. We sought to determine the effect of tacrolimus and sirolimus on mitochondrial respiration in endothelial cells. Interestingly, we found that sirolimus and tacrolimus treatments had no significant effect on mitochondrial respiration in endothelial cells suggesting the mechanism by which the abnormalities observed in endothelial function is not driven by metabolic alterations **(Figure 5)**.

**Fig 5.**
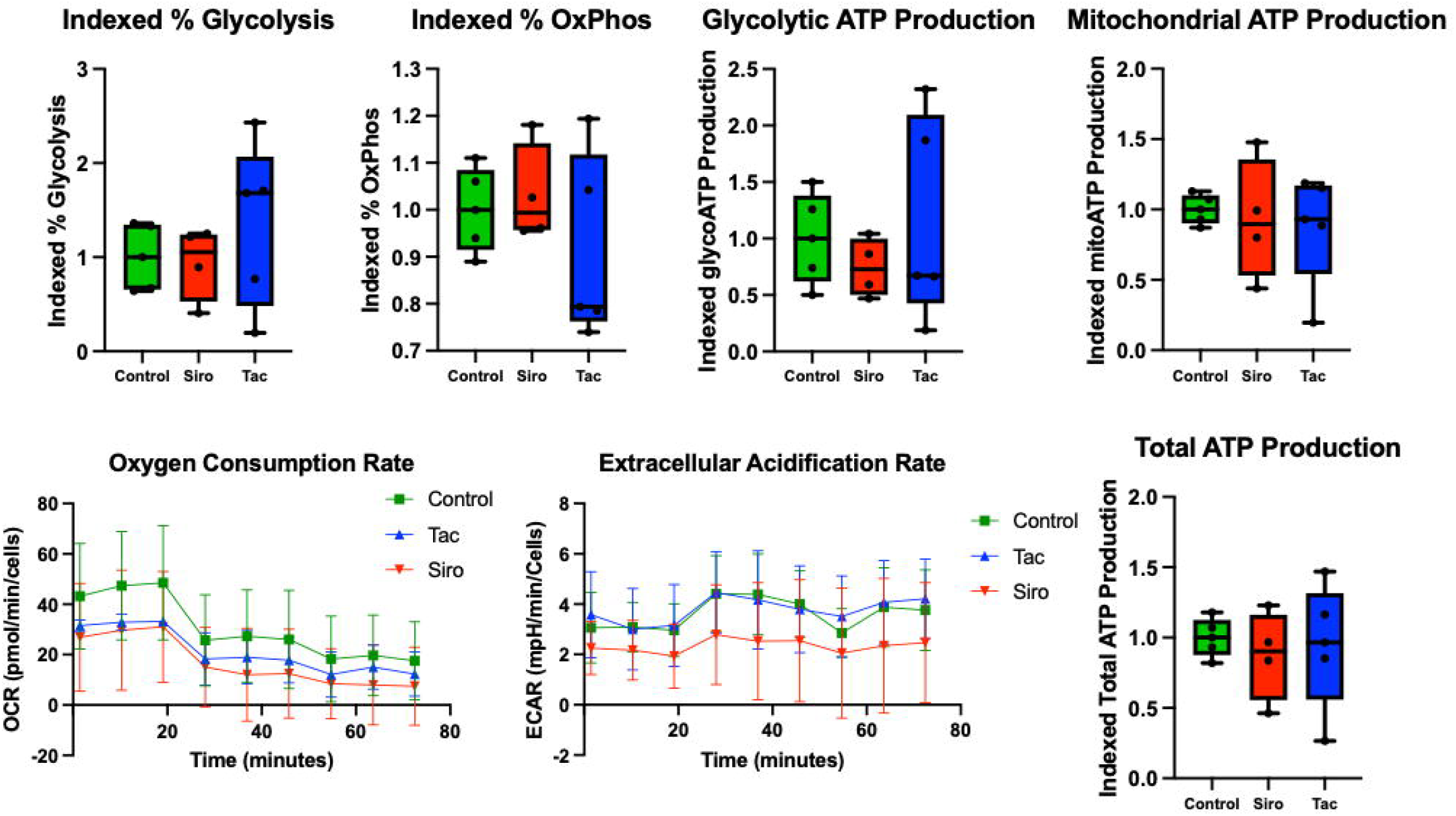
Tacrolimus and sirolimus treatment of iPSC-ECs does not affect mitochondrial respiration. iPSC-ECs had no difference in glycolytic ATP production, mitochondrial ATP production, total ATP production, or the relative contribution of glycolysis and oxidative phosphorylation (OxPhos) to ATP production.

## Discussion

Immunosuppressant use is associated with vascular outcomes, with differential rates of hypertension, dyslipidemia, and atherosclerosis in patients treated with mTOR inhibitors and calcineurin inhibitors^3^. Given the systemic effects of the drugs and the multi-organ interactions implicated in vascular function, delineating cell-specific alterations is difficult in vivo. Here, we show that sirolimus and tacrolimus have direct effects on endothelial cell function. Specifically, sirolimus treatment led to a decrease in cell migration, proliferation, and acetylated LDL uptake, while both tacrolimus and sirolimus reduced nitric oxide release from iPSC-ECs. These effects may provide mechanistic insight into the vascular phenotype associated with mTOR inhibitor and calcineurin inhibitor use.

Hypertension is a well-described side effect of immunosuppressive medication use, particularly calcineurin inhibitors, and is associated with an increased risk of coronary artery disease, cerebrovascular events, renal dysfunction, and adverse cardiovascular remodeling^22^. Cyclosporine A and tacrolimus are associated with promoting direct vasoconstriction by increased tone of vascular smooth muscle and activation of endothelin-1 receptor^23–25^. mTOR Inhibitors, on the other hand, have been associated with a lower risk of hypertension compared to calcineurin inhibitors when used in solid organ transplant recipients^26^. Sirolimus is thought to prevent endothelial hyperplasia and dysfunction^27^. Here, we found that in iPSC-ECs, treatment with sirolimus and tacrolimus decreased nitric oxide production. The reduced nitric oxide in response to tacrolimus is in line with other models of calcineurin inhibitors and observed increased hypertension. Our sirolimus results are in line with clinical observations of endothelial vasomotor dysfunction in response to sirolimus exposure^28^. Our findings support both immunosuppression drugs reducing endothelial nitric oxide production.

Immunosuppressive medications are associated with dyslipidemia^13,29^. Each drug class is associated with individual variations in affected lipid particles and more importantly in the conferred risk of atherosclerosis. Among calcineurin inhibitors, cyclosporine A is associated with a dose-dependent increase in total cholesterol and LDL cholesterol, a decrease in high-density lipoprotein (HDL) cholesterol, and an increase in serum triglycerides^30^. Tacrolimus use is associated with a milder dyslipidemia profile^31^. mTOR inhibitors, particularly sirolimus, are stronger inducers of hyperlipidemia than calcineurin inhibitors, associated clinically with an increase in serum LDL and triglyceride levels^32^. The mechanism remains unclear, although it may be due to a combination of reduced catabolism, an increase in the free fatty acid pool, increased hepatic production of triglycerides, and secretion of very low density lipoprotein^33,34^. Interestingly, despite the increase in serum lipids, mTOR inhibitors are associated with an overall lower risk of atherosclerosis^6^. Here, we show that sirolimus treatment of endothelial cells decreases LDL uptake, while tacrolimus treatment maintains ability to uptake LDL. This may in part explain the differential effect of mTOR inhibitors on dyslipidemia and atherosclerosis.

Immunosuppressive agents directly contribute to abnormal vascular remodeling that may drive cardiovascular adverse events, independent of hypertension or dyslipidemia^7^. Defining this risk and the contributing mechanisms for each drug is important in order to ensure appropriate follow up and identify potential actionable targets to modify the risk profile. Calcineurin inhibitors have been associated with increased risk of allograft vasculopathy, a notable complication of transplanted hearts which represents a major driver of graft dysfunction and has significant implications for quality of life and longevity of heart transplant recipients^35^. In animals treated with tacrolimus, adverse remodeling features described include vascular stiffness, thickening, inflammation, and fibrosis^36,37^. Proposed mechanisms include decreased fibrinolytic activity in vessel walls, and possibly increased intracellular calcium in vascular smooth muscle cells^38–40^. Both sirolimus and everolimus have been associated with a more favorable vascular profile with clinical efficacy in reducing the rate of progression of cardiac allograft vasculopathy, which has led to widespread use in heart transplant recipients^41,42^. mTOR inhibitors treatment has been shown to minimize intimal hyperplasia, vascular smooth muscle proliferation, and infiltration by inflammatory cells^9,43,44^. Here, we show that sirolimus treatment of endothelial cells decreases their proliferation, migration, and nitric oxide release. While this may translate to an increased risk of capillary leak and lymphedema; this may confer a protective effect in terms of coronary pathology such as atherosclerosis and CAV.

We show distinct endothelial function profiles in response to tacrolimus or sirolimus treatment. Sirolimus exerted a broader effect on endothelial cell function, while tacrolimus primarily affected nitric oxide release. The knowledge of these endothelial-specific effects can be used to inform further studies on immunosuppression associated vascular remodeling in multicellular or organ models.

## Statements

### Statement of Ethics

#### Study approval statement

The iPSC lines used in this study were obtained from the Stanford Cardiovascular Institute Biobank (https://med.stanford.edu/scvibiobank.html). The recruitment and reprogramming of the iPSC lines was done with written informed consent with Stanford IRB Approval (Protocol 29904)

## Author Contributions

AE: performed experiments, analyzed data and writing of manuscript

RD: performed experiments, analyzed data, assisted in manuscript writing and figures

DW: performed experiments and analyzed data

SC: performed experiments and analyzed data

IYC: experimental design, analysis of the data and writing of manuscript

NS: experimental design, analysis of the data and writing of manuscript

KS: conceptualization of the project, experimental design, analysis of data, and writing of the manuscript

## Data Availability Statement

The data generated and analyzed during the current study are available from the corresponding author on reasonable request. The data that support the findings are not publicly available because they make compromise privacy of original donor but are available from KS.

